# Age-Invariant Genes: Multi-Tissue Identification and Characterization of Murine Reference Genes

**DOI:** 10.1101/2024.04.09.588721

**Authors:** John T. González, Kyra Thrush, Margarita Meer, Morgan E. Levine, Albert T. Higgins-Chen

## Abstract

Studies of the aging transcriptome focus on genes that change with age. But what can we learn from age-invariant genes—those that remain unchanged throughout the aging process? These genes also have a practical application: they serve as reference genes (often called housekeeping genes) in expression studies. Reference genes have mostly been identified and validated in young organisms, and no systematic investigation has been done across the lifespan. Here, we build upon a common pipeline for identifying reference genes in RNA-seq datasets to identify age-invariant genes across seventeen C57BL/6 mouse tissues (brain, lung, bone marrow, muscle, white blood cells, heart, small intestine, kidney, liver, pancreas, skin, brown, gonadal, marrow, and subcutaneous adipose tissue) spanning 1 to 21+ months of age. We identify 9 pan-tissue age-invariant genes and many tissue-specific age-invariant genes. These genes are stable across the lifespan and are validated in independent bulk RNA-seq datasets and RT-qPCR. We find age-invariant genes have shorter transcripts on average and are enriched for CpG islands. Interestingly, pathway enrichment analysis for age-invariant genes identifies an overrepresentation of molecular functions associated with some, but not all, hallmarks of aging. Thus, though hallmarks of aging typically involve changes in cell maintenance mechanisms, select genes associated with these hallmarks resist fluctuations in expression with age. Finally, our analysis concludes no classical reference gene is appropriate for aging studies in all tissues. Instead, we provide tissue-specific and pan-tissue genes for assays utilizing reference gene normalization (i.e., RT-qPCR) that can be applied to animals across the lifespan.

## Introduction

Aging, the accumulation of cellular, molecular, and physiological alterations in an organism over time, increases the risk of dysfunction, chronic disease, and mortality [1]. The advent of next-generation sequencing and other high-throughput technologies has allowed for data-driven analyses to discover age-linked gene expression changes and dysregulation. However, little effort has been directed toward identifying and understanding **age-invariant genes** – those that remain unchanged throughout the aging process. The utility of such genes would be twofold: 1) they can be used as reference genes in quantitative assays, and 2) they may share molecular features that allow them to resist changes with age.

The transcriptome has been shown to exhibit substantial remodeling during the aging process, and there is evidence that many of these changes may drive declines in cellular function. By employing bulk RNA-seq across 17 mouse tissues, Schaum et al. identified clusters of genes with similar age trajectories associated with the hallmarks of aging [2]. Gene clusters increasing in expression included immune and stress response genes, while those decreasing in expression included genes involved in the extracellular matrix, mitochondria, and protein folding [2]. Overall, a global decrease in gene expression has been reported to occur with aging, such that when comparing older animals to younger animals, differentially expressed genes tend towards downregulation [3]. For tissue-specific genes, a divergence or specialization of distinct cell types is observed during development, whereas aging has been associated with a loss of specificity in transcriptional profiles [4] and an increase in transcriptional noise (increased variance between individuals) [5–7]. Interestingly, genes subject to age-related change have been linked to specific features, including transcript length and association with CpG islands [8,9].

Studying age-invariant genes that do not change their expression and remain stable throughout the aging process may uncover complementary aging mechanisms. The notion of invariant genes has been a focus of biomedical research for over 50 years, but their study has been confined to young organisms or cell line perturbations [10]. Due to their relative stability, invariant genes have been utilized as internal reference controls for gene expression assays. Initially coined as housekeeping genes, these invariant genes are constitutively expressed at high levels, are subject to low fluctuations, and are often essential for proper cellular function [10–12]. The changing definition of the term “housekeeping gene” led the *Minimum Information for Publication of Quantitative Real-Time PCR Experiments* (MIQE) guidelines to update the term used for normalization to **reference genes (RGs)** [13], and we will utilize this term. There is no absolute standard list of RGs; many classical RGs, including glyceraldehyde-3-phosphate dehydrogenase (GAPDH), actin β (ACTB), and β2-microglobulin (B2M), were found to be highly variable in certain contexts [11,14]. Although an ultimate RG may not exist (consistent across all possible tissues, cell types, cell cycle stages, experimental conditions, and developmental phases), identification of invariant genes in specific contexts and sample types is possible [14,15].

Little work has been done to identify and validate RGs that are stable throughout the aging process, i.e., age-invariant reference genes. These genes would be invariant across the lifespan, either within any given tissue (tissue-specific) or across all tissues (pan-tissue). Aging is known to impact classical reference gene expression: a mouse study, for example, found age, sex, and frailty explicitly alter the expression of a majority of classical RGs examined [16].[11,14] [29]. Within the aging field, studies are restricted to RGs identified in other fields rather than using a novel, aging-focused analysis. The few available studies examining RGs in aging employ targeted RT-qPCR validation of some of the aforementioned classical transcripts and recommend different RGs based on the genes and the parameters included. For example, GUSB increased with age in mouse skeletal muscle, making it a poor RG in that context, but it was the best RG candidate in human peripheral blood mononuclear cells [16–20]. Another salient example for aging is Cdkn1a/p21. Cdkn1a/p21 is often utilized as a reference gene in RT-qPCR normalization literature [20], even though it simultaneously serves as a marker of cellular senescence–one of the major hallmarks of aging, which is defined by change over time [21,22]. Thus there is a pressing need to identify RGs appropriate for aging studies.

We now have the tools and datasets to identify age-invariant RGs. The first iterations of reference genes, which compose a majority of popular RGs, were not experimentally determined but selected because they were detected in all tissues and assumed to have little variability [10,23]. With the development of 21st-century microarray and next-generation sequencing technologies, this question can finally be tackled from a data-rich perspective [23]. RNA-seq datasets have been successfully used to experimentally identify RGs in healthy human tissue [10,11], mammalian animal models [14,24], non-mammalian organisms [25], disease conditions [26] and even single-cell populations [27]. The variables included in the datasets for these analyses determine the application constraints of the resulting RGs. Novel data-rich unsupervised techniques paired with next-generation sequencing data remain an untapped resource for identifying RGs for aging studies and more fully understanding the dynamics of transcriptional change (or lack thereof) with aging.

Here, we leverage published approaches for RG identification [10] with appropriate refinements (**Figure 1A**) and apply them to public bulk RNA-seq datasets with samples collected across the full lifespan (**Figure 1B**) to identify age-invariant genes. We show that, unlike our age-invariant genes, no classical RG is suitable for aging studies across all tissues (**Figure 1C**); and characterize features and functions of these age-invariant genes (**Figure 1D**). Of note, we opted to focus on the subset of age-invariant genes that can also serve as RGs – those that are also relatively highly expressed – due to their practical applications.

**Figure 1:**
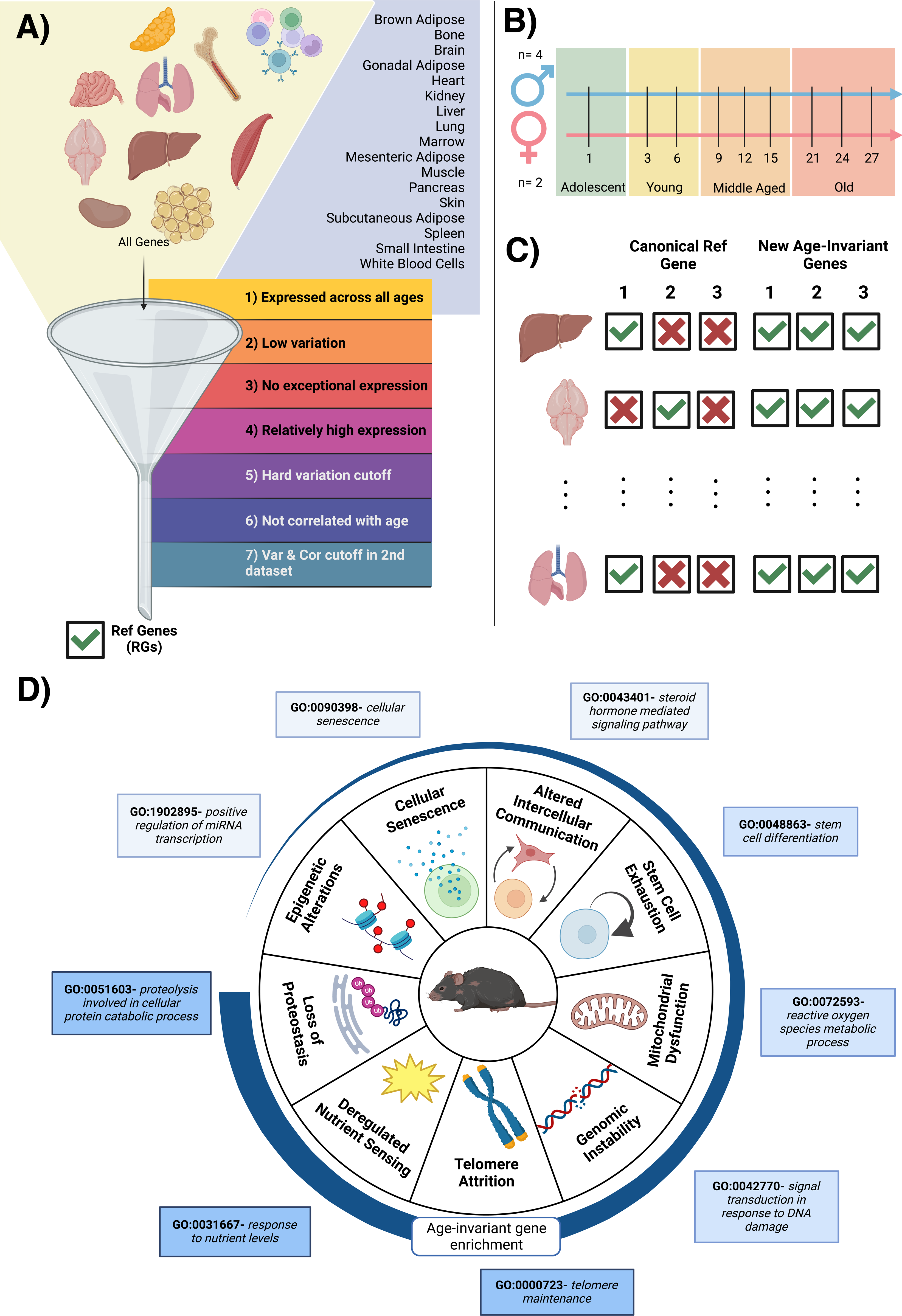
Visual Diagram of Article Contents. A) Bulk RNA-seq data from 17 murine tissues (GSE132040) were sequentially filtered through 7 criteria. Steps 1-4 are adapted from previous publications. We added criteria filters 5 and 6 to ensure low variation and no correlation with age. Criteria filter 7 was validation of low variation and no age correlation, performed in a second dataset for 11 of the 17 tissues. B) Sample gender, age and life stage distributions of the samples in the dataset. A full table of samples can be found in Supplementary Table 10. C) Canonical reference genes are not applicable to all tissues in an aging context but age-invariant genes introduced here are. D) Tissue aging-invariant genes are enriched to different extents for gene ontology terms associated with hallmarks of aging. Age-invariant genes have low enrichment in some (e.g. epigenetic alterations GO terms) and high enrichment in others (e.g. loss of proteostasis GO terms). Created with BioRender.com

## Results

### Identification of candidate age-invariant genes from RNA-Seq data

Bulk RNA-seq data from the Tabula Muris Senis study [2] were utilized for age-invariant RG discovery. We analyzed 17 tissues: brown adipose tissue (BAT), bone, brain, gonadal adipose tissue (GAT), heart, kidney, limb, liver, lung, marrow, mesenteric adipose tissue (MAT), pancreas, subcutaneous adipose tissue (SCAT), skin, small intestine, spleen and white blood cells (WBCs). We performed quality control and only utilized samples where we could verify the tissue label (**Supplementary** Figure 1**; Methods**). The dataset contained female and male mice representing the 4 major lifespan stages: adolescent (1mo), young (3 and 6mo), middle-aged (9, 12, and 15mo), and old (21, 24, and 27mo) [28] (**Figure 1A-B**).

Tissues were independently analyzed by sequentially applying 7 filtering criteria through each tissue’s gene set (**Figure 1A**). Here, we utilize expression counts normalized to **Transcripts Per Million (TPM)** [29], which is similar to RT-qPCR as it approximates relative molar RNA concentration, as well as **Trimmed Mean of M (TMM)** [30], which leverages inter-sample information to reduce sensitivity to gene outliers. Both normalization techniques performed similarly well at identifying RGs in a recent systematic comparison of normalization methods [26]. Our approach leverages two different normalization techniques to reduce artifacts specific to individual methods. Each criterion, or filter, was applied to each tissue individually with both normalization methods; genes were only included in the tissue-filter gene list if they satisfied the requirement in both TPM and TMM normalized datasets.

Our filtering criteria are listed below. The filtering pipeline was applied to each tissue separately, with samples spanning the lifespan stages defined in **Figure 1B**. Although some genes have been identified as age-invariant within multiple tissues, this does not suggest they are invariant to tissue type and thus should still be applied in a tissue-specific manner. Criteria 1-4 are adapted from an approach frequently used for RG identification from RNA-seq data [10,25]:

1. <u>Continuous expression:</u> Non-zero expression in all samples.
2. <u>Low variance:</u> The **standard deviation (SD)** of the log2 normalized gene (x) expression for all samples (i) is less than 1. V, (_2_()) < 1
3. <u>No exceptional expression/outliers:</u> log2 normalized values are within two units of the gene’s mean(removing genes with data points four-fold away from the gene mean). V, | _2_()-(_2_()) | < 2
4. <u>Medium to high gene expression:</u> the gene’s log2 normalized expression mean is above the mean of all the genes expressed in the particular tissue V, (_2_()) > (_2_()) To ensure age-invariant gene list quality, we added two new filters to the identification criteria:
5. <u>Low</u> **<u>coefficient of variation (CV)</u>**: The percent coefficient of variation (%CV), the ratio of the standard deviation to the mean, is lower than 20%. V, (_2_()) / (_2_()) < 20
6. <u>No correlation between gene expression and age</u>: Gene expression correlation with age is not statistically significant (no p-value under 0.05) Finally, we performed external validation:
7. Filters 5 and 6 were applied in publicly available validation datasets with bulk tissue RNA-seq data from mice. Tissues with a validation dataset were BAT, brain, heart, kidney, muscle, liver, lung, SCAT, skin, small intestine, and WBCs (11).

The filters progressively refined the list of both tissue-specific (**Figure 2A, Supplementary Table 1**) and pan-tissue age-invariant genes (**Figure 2B**, **Table 1**). For reference, **Supplementary Table 2** lists information on each gene’s %CV, slope with age, and correlation with age in each tissue, allowing readers to select their own cutoffs if they choose. **Supplementary Tables 3-9** contain lists of all genes that passed each consecutive filter in each tissue.

**Figure 2:**
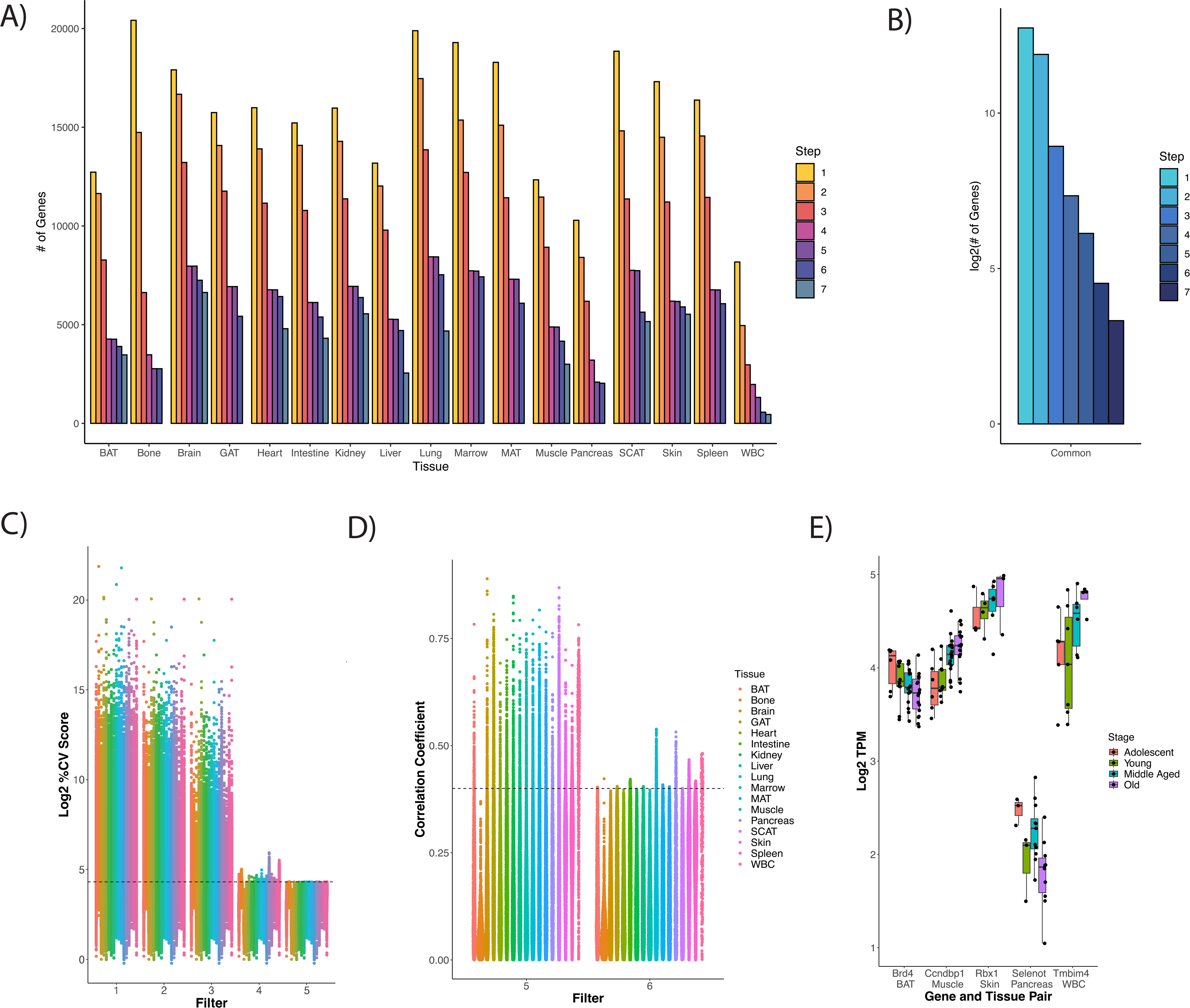
Gene Selection Process and Rationale. A) Gene count number remaining after each criteria/filter step for each tissue. B) Gene count present across all tissues at each step, presented on a log2 scale. C) % Coefficient of Variance (CV) for each gene calculated as SD/mean*100 distribution of log2 TPM gene expression values. Genes that satisfy every subsequent filter are plotted by the last filter applied. Filters 1-3 slowly decrease %CV and the cumulative effect of filters 1-4 generally results in a %CV of approximately 20%. Filter 5 imposes a strict %CV < 20% requirement for all tissue-gene pairs. D) Age information must be included in exclusion criteria as low variation genes can still have a high correlation with age. Filter 6 (Spearman correlation p-value based removal) removes highly age-correlated genes. Dashed line corresponds to a correlation coefficient (y-axis) of 0.4, which for most tissues corresponds to a significant correlation with p = 0.05. Exact CV and age correlation information is found in Supplementary Table 2, in case readers wish to utilize other cutoffs in selecting RGs. E) Log2 TPM (y-axis) values by life stage (color) for specific gene-tissue pairs (x-axis) for genes that satisfy filters 1-5, but are eliminated by filter 6. Boxplot line represents the group median while lower and upper limits of the boxplot correspond to the first (25%) and third (75%) quartiles.

**Table 1:**
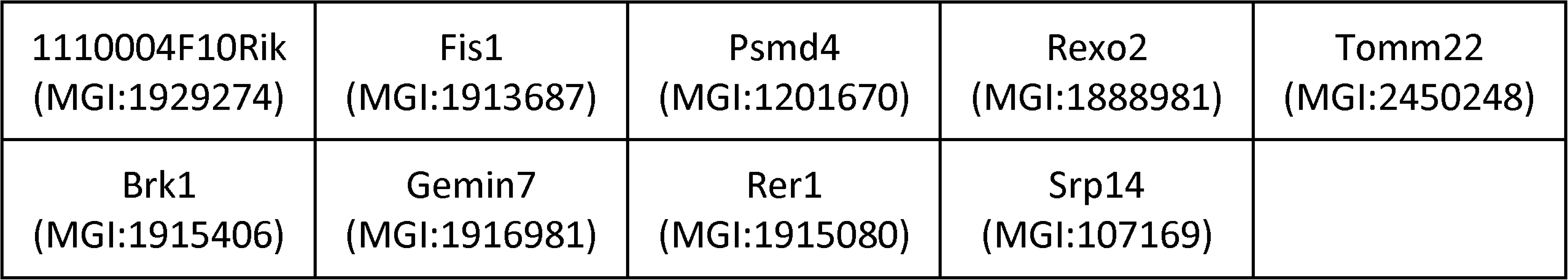
MGI symbol and ID for our 9 pan-tissue age-invariant genes. These genes were present across all tissues after all filtering steps and validation.

|  |  |  |  |  |
| --- | --- | --- | --- | --- |
| 1110004F10Rik<br>(MGI:1929274) | Fis1<br>(MGI:1913687) | Psmd4<br>(MGI:1201670) | Rexo2<br>(MGI:1888981) | Tomm22<br>(MGI:2450248) |
| Brk1<br>(MGI:1915406) | Gemin7<br>(MGI:1916981) | Rer1<br>(MGI:1915080) | Srp14<br>(MGI:107169) |  |

There were a few notable modifications to the original pipeline. First, we modified Criterion 4, which selects for relatively highly expressed genes and, therefore, is easily detected by RT-qPCR [10]. Because each tissue had different gene count distributions **(Supplementary** Figure 2A), we deviated from the previous use of an arbitrary cutoff and employed an adjusted cutoff, removing genes with means below the mean of all genes expressed in a given tissue (log2 transformed) [25]. Consistent with previous publications [25], the cumulative effect of filters 2 (standard deviation cut-off) and 4 (mean cut-off) resulted in a percent coefficient of variation (%CV) of about 20% in most tissues (**Figure 2C**). However, given the lower average normalized gene expression in some tissues (Bone, Pancreas, Spleen WBC), genes in these tissues surpassed this threshold. To ensure the genes obtained were truly low variance, we applied a hard cut-off of 20% CV (Filter 5). This approach combines Eisenberg et al.’s low variance definition of RGs and their alternative approach: mid-to-high expression [10].

Second, we added Filter 6 to ensure age invariance. We had initially hypothesized that simply analyzing samples with a wide age range using the typical RG pipeline (filters 1-4) would be sufficient to filter out genes that change with age. Indeed, adding age groups to the analysis progressively discarded genes during the filtering process (**Supplementary** Figure 3A). This, however, could be due to the increase in samples (n) included in the analysis. To test whether the wide age range alone contributes important information, we applied the steps of the standard pipeline (filters 1-4) on samples belonging to only a particular lifespan stage and compared it to a cross-stage control with the same n. Including a wide range of ages by using cross-stage analysis discarded more genes compared to single-stage analysis for adolescent, middle-aged, and old stages (**Supplementary** Figure 3B). Surprisingly, this was not the case for the young adult stage (3-6mo old); we found this was likely due to a subset of genes that have high expression variability in young adults but are stable in other life stages (**Supplementary** Figure 3C-F). Regardless of lifestage analyzed, this pattern held true. Genes identified as age-invariant (with filters 1-4) only in young samples (**Supplementary** Figure 3C); in young samples and other lifestages (analyzed separately)(**Supplementary** Figure 3D); in lifestages except young (**Supplementary** Figure 3E); or with the full dataset, i.e., all lifespan stages, (**Supplementary** Figure 3F) reveal a similar pattern: some genes have higher variance (%CV) in young and old populations. This is reflected by the rightward shift in young and old samples. Young samples have an overall higher proportion of high variance (over the 20%CV dotted line) genes than old ones (**Supplementary** Figure 3C-F). Thus, simply utilizing a wide age range in the typical pipeline does not necessarily help identify age-invariant genes. Furthermore, we found that some genes obtained through filters 1-5 still changed with age (**Figure 2 D-E**). To address this finding, we added criterion 6, removing genes with statistically significant correlations with age for each tissue (**Figure 2D**).

Finally, to decrease the number of false positives, we validated the gene lists using a second bulk mRNA-seq dataset for 11 out of 17 tissues (except for bone, GAT, marrow, MAT, pancreas, and spleen). The number of validated genes is displayed in **Figure 2A-B** as Step 7. Specific counts and percentages can be found in **Supplementary Table 1**. For nearly all tissues, a supermajority (>70%) of candidate age-invariant genes were validated, except in the liver (54%) and lung (62%). The fewest number of age-invariant genes was observed in WBCs, possibly due to large changes in distributions of cell types over shorter timescales [31,32](**Figure 2A**).

### RT-qPCR validation of Novel age-invariant reference genes

Our analysis identified many tissue-specific age-invariant RGs (**Supplementary Table 9**), as well as 9 such genes common to all tissues (pan-tissue). Some classical RGs are not age-invariant genes (**Figure 3A-C**). In fact, no classical RGs were age-invariant across every tissue (**Figure 3A**). Thus, we propose a new list of 9 age-invariant genes common to all 17 tissues that can be used in studies with aged animals (**Table 1**).

**Figure 3:**
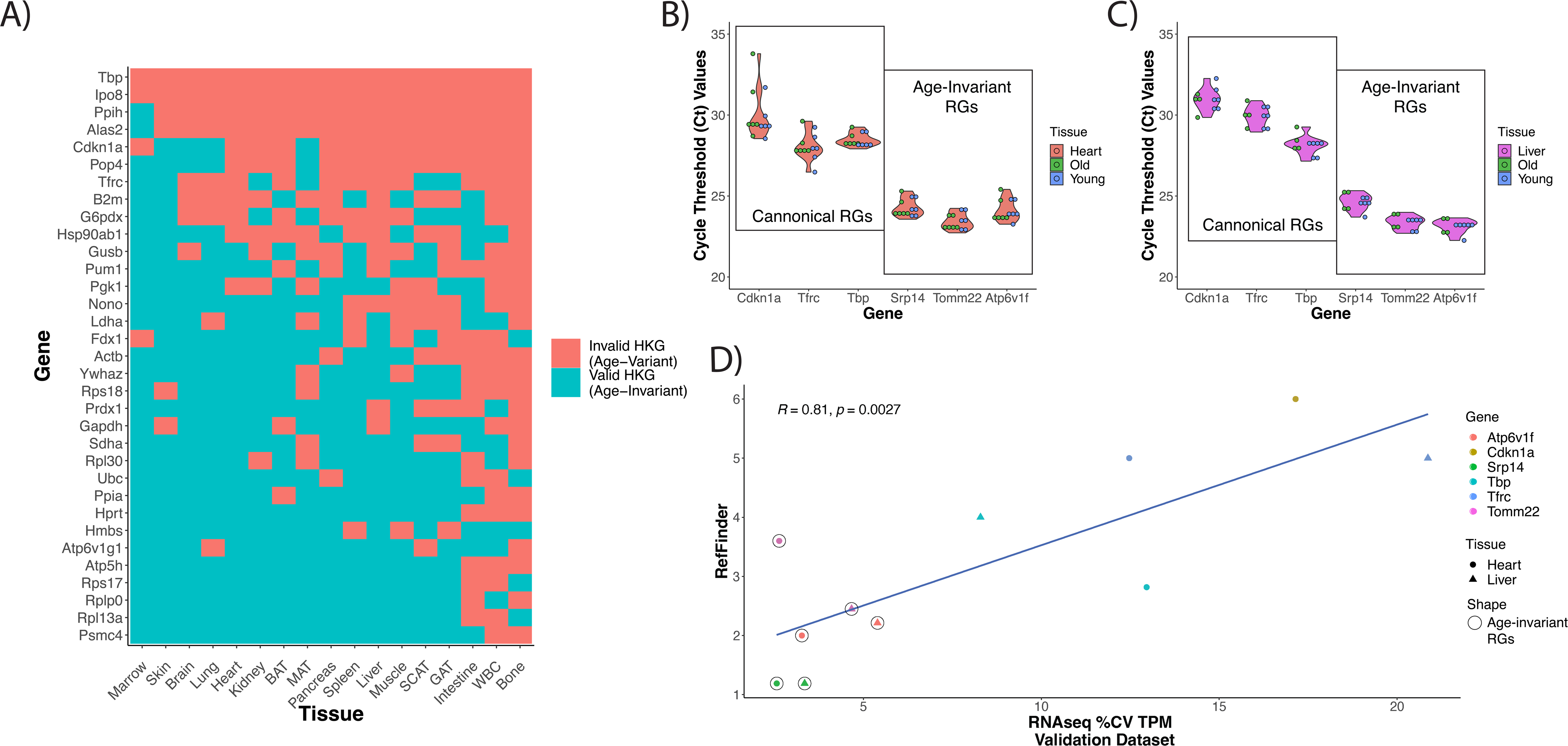
Classical & Novel RG Performance in Aging Samples. A) Aging RG status of classical reference gene by tissue. Genes that are age-invariant and therefore valid RGs are depicted in blue while their age-variant counterparts, which were not present in the gene list after filtering, appear in red. B-C) Individual gene cycle threshold (Ct) results from validation RT-qPCR tissues in heart (B) and liver (C) for selected classical RGs and novel age-invariant RGs. D) RT-qPCR Gene RefFinder score and mRNA-seq %CV in heart and liver. Age-invariant genes are distinct from and outperform canonical RGs in %CV (Welch Two Sample t-test p-value= 0.006553) and RefFinder qPCR scores(Welch Two Sample t-test p-value = 0.02401). RefFinder and %CV scores were calculated from in-house and public validation datasets respectively. RefFinder score was based on BestKeeper, NormFinder, GeNorm and comparative delta-Ct values. Circled points indicate novel age-invariant RGs (Two pan-tissue: Tomm22 and Srp14; and one heart and liver age-invariant gene: Atp6v1f) while uncircled points specify classical RGs from Figure 3A.

These RG sets can be utilized in the context of northern blot, RT-qPCR, and some RNA-seq normalization strategies in aging studies. Researchers have the choice of selecting from a tissue-specific gene list or from the nine pan-tissue genes. To validate this, independent samples were used to generate RT-qPCR data for three age-invariant genes identified by our computational pipeline: Atp6v1f, Srp14, and Tomm22 (**Figure 3B-D**). Atp6v1f is an age-invariant gene shared by the two tissues assayed: the liver and heart. The other two are pan-tissue age-invariant genes. The novel samples consisted of mouse heart and liver samples in four categories: old (∼19mo) female, old male, young (∼8mo old) female, and young male. We compared these against three classical RGs: Cdkn1a, Tbp, and Tfrc. Classical reference genes generally had a wider cycle threshold distribution than the age-invariant genes, with Tbp being the most stable among them, followed by Tfrc and Cdkn1a (**Figure 3B-C**). Cdkn1a codes for cyclin-dependent kinase inhibitor 1A, also known as p21. Given that Cdkn1a is widely used as a marker of cell senescence [22], it is not surprising that it has a high degree of variability despite it being widely considered an RG in RT-qPCR normalization literature [17].

To assess gene RT-qPCR stability in the context of aging, we calculated the expression stability across multiple algorithms: BestKeeper [33] (**Supplementary** Figure 4A-B), geNorm [34] (**Supplementary** Figure 4C-D), NormFinder [35] (**Supplementary** Figure 4E-F), and delta-CT method [36] (**Supplementary** Figure 4G-H). These scores were utilized to calculate the summary RefFinder score (**Figure 3D, Supplementary** Figure 4I) [37]. TPM %CV for the discovery (**Supplementary** Figure 4A**, C, E, G, J)** and validation **(Figure 3D, Supplementary** Figure 4B**, D, F, H)** RNA-seq datasets strongly correlate with all stability algorithm values calculated on our in-house samples. **Figure 3D** displays the correlation between the TPM %CV values of the external validation dataset and the RefFinder score of our in-house validation samples (Pearson correlation = 0.81, p-value = 0.0027). This suggests that %CV from normalized RNA-seq samples could be used as an indicator of candidate reference genes for RT-qPCR experiments subject to the same conditions. By both metrics, the newly identified age-invariant genes outperformed the classical RGs: these RGs are statistically different in both %CV (Welch Two Sample t-test p-value= 0.006553) and RefFinder qPCR scores (Welch Two Sample t-test p-value = 0.02401) This suggests age-invariant genes common across all tissues (Srp14 and Tomm22) or particular tissues (Atp6v1f in heart and liver) could be applied as part of normalization in age-related transcriptomic research. A combination of more than one of the age-invariant genes is recommended for RT-qPCR experiments, per the MIQE guidelines [13].

### Overlapping pathways for aging stable and aging dysregulated genes

Gene enrichment analysis of the tissue-specific age-invariant genes revealed a large number of statistically significant GO biological pathway terms **(Supplementary** Figure 5). As expected, the most enriched terms were largely involved in basic metabolic and structural processes **(Figure 4A)**. We also noted many enriched terms were related to the hallmarks of aging [38], which was surprising considering that hallmarks of aging are typically thought to involve processes that change with age. As an initial step to systematically assess the presence of stably transcribed genes in these hallmarks, we compared the enrichment scores of our tissue age-invariant gene lists with previously published enrichment terms associated with age dysregulation and disease [2,39].

**Figure 4:**
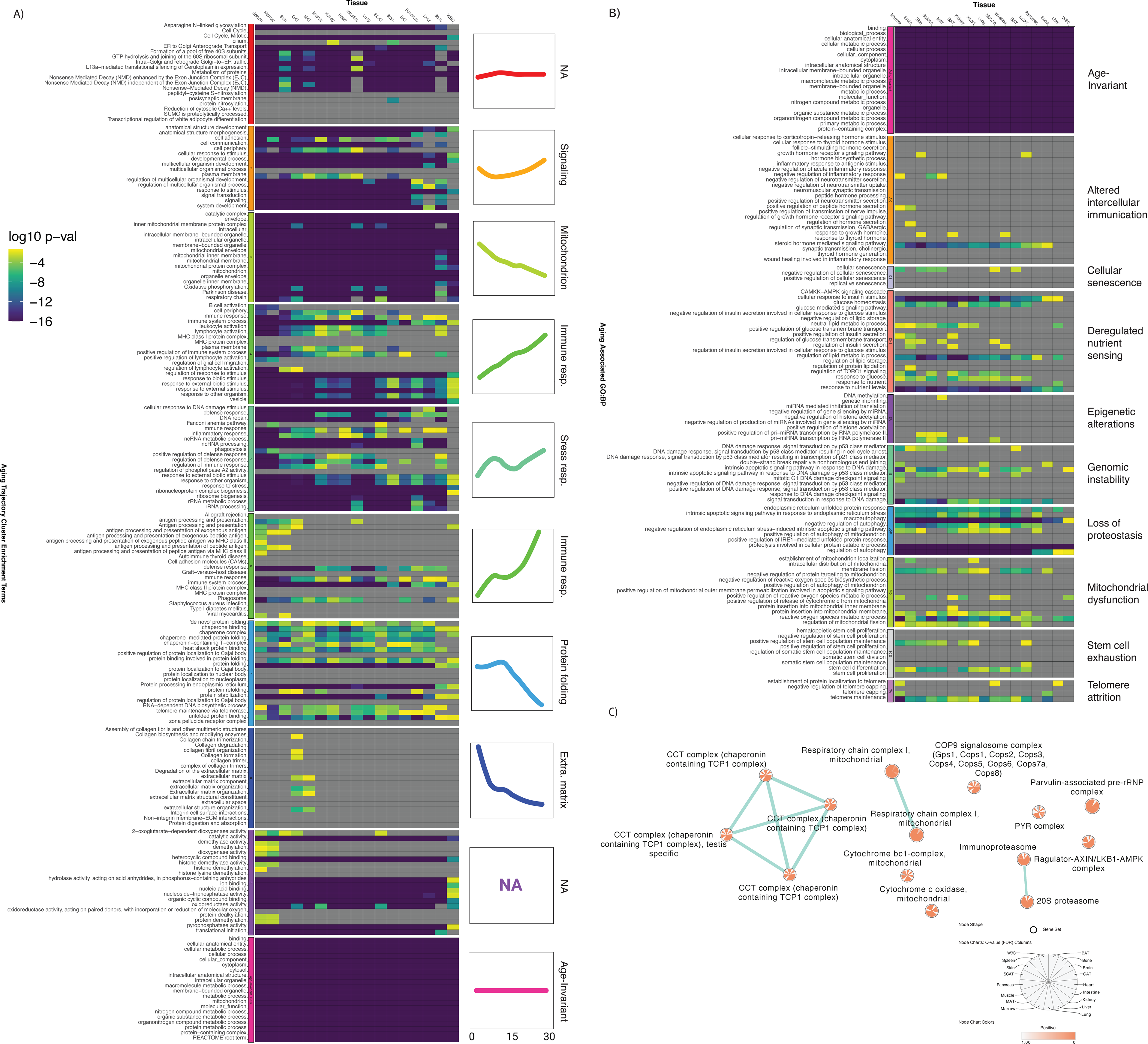
Age-Invariant Genes are Enriched for Dysregulated and Aging Disease Associated Gene Functions. A) Tissue age-invariant genes are enriched for some GO, KEGG and REACTOME terms associated with linear and non-linear aging trajectories. Left labels correspond to enrichment terms originally classified in 9 trajectory groups. Age-invariant labels at the very bottom (pink) refer to genes identified in this paper. Heatmap columns correspond to different tissues, while rows correspond to different terms. B) Age-invariant genes are enriched for GO Biological Processes associated with age-related disease in humans. C) Tissue age-invariant genes are enriched for certain protein complexes. Gene lists are enriched for CCT complex, electron transport chain (respiratory complex I and cytochrome C), proteasome, Cop9 signalosome, PYR, Parvulin-associated pre-rRNP and Regulator-AXIN/LKB1-AMPK complexes in CORUM analysis.

We first compared our enrichment scores with the top terms associated with mouse transcriptome aging clusters, each displaying a different trajectory with aging **(Figure 4A)**. The top enrichments of these 10 clusters, obtained from the same dataset we performed our discovery on, are associated with hallmarks of aging like protein folding, inflammation, and mitochondrial function [2]. We found our tissue gene sets were significantly enriched in many, but not all, of the clusters. Of note is cluster 3, linked to mitochondrial dysfunction, where age-invariant genes are highly enriched for every term of this cluster. Age-invariant genes are also heavily represented in stress response (cluster 5), signaling (cluster 2), and protein stability (cluster 7). Interestingly, within the protein stability cluster, age-invariant genes were enriched in terms involved in protein folding, processing, and stabilization but not in terms involved in protein localization. The clusters with the least age-invariant genes were those associated with immune response and extracellular matrix. This suggests that hallmarks themselves, or mechanisms within an aging hallmark, can be separated by the presence or absence of age-invariant genes.

Cluster 1 from Schaum et al. is defined as genes that do not change with age and, as expected, has a large overlap with our tissue age-invariant gene sets. Cluster 1 was defined by having the least amplitude (change with age) and least variability. Interestingly, throughout the 17 tissues, only ∼33-40% of our age-invariant genes were in Schaum et al.’s cluster 1. The genes not shared between both methods likely reflect the difference between relatively a stable group of genes identified by hierarchical clustering and individual age-invariant genes identified due to their characteristics (as well as our RG requirement that genes be highly expressed) [2,40]. In RNA-seq, genes with low expression demonstrate significant technical noise making it difficult to assess true biological variability related to age or other factors, and are often filtered out of differential expression studies [41], so our requirement for high expression is useful for focusing on age-invariant genes.

The other ontology terms we examined came from an analysis of age-related diseases and aging hallmarks **(Figure 4B)**. Unlike Schaum et al., who used a completely unsupervised approach, Fraser et al. used genes associated with human age-related diseases in a genome-wide association study to define GO biological pathways related to both disease and at least one aging hallmark [39]. Most hallmarks have at least one GO term enriched for age-invariant genes across most of the tissues analyzed (e.g., “steroid hormone-mediated signaling pathway” in altered intercellular communication; “cellular response to insulin stimulus” and “response to nutrient levels” in deregulated nutrient sensing; “macroautophagy” and “regulation of autophagy” for loss of proteostasis; “reactive oxygen species metabolic process” in mitochondrial dysfunction; and “telomere maintenance” in telomere attrition). On the other hand, virtually no GO term related to cellular senescence and epigenetic alterations had high proportions of stably transcribed genes. According to this alternative way of identifying gene ontology terms associated with aging hallmarks, age-invariant genes continue to be enriched in these terms.

To better understand the implications of some of these stable pathways, we used the comprehensive resource of mammalian protein complexes (CORUM) database to perform enrichment analysis **(Figure 4C)**[42]. The enriched complexes are consistent with our enrichment results in this data thus far. Complexes involved in mitochondrial function (respiratory chain complex I and cytochrome c oxidase), stress response & signaling (Regulator-AXIN/LKB1-AMPK complexes), and protein stability (COP9 signalosome, proteasome, Parvulin-associated pre-rRNP, and Chaperonin containing TCP1 Complex) are enriched in age-invariant genes.

Our analyses reveal multiple age-invariant genes within pathways that are either dysregulated with aging **(Figure 4A, C)** or associated with aging pathologies **(Figure 4B)**. Pathways related to the extracellular matrix, cellular senescence, and epigenetic alterations seem particularly devoid of stably expressed genes. These findings are not due to the high expression requirement for our age-invariant genes, as removing this requirement produced similar results **(Supplementary** Figure 6).

### Age-invariant gene features

Features of genes that change with age have long been a point of discussion in aging transcriptome research, but little is known about the genes that are able to withstand the effects of time. We tested whether our genes have the opposite features to those described in age-dysregulated transcriptome analyses. The features examined are CpG content, DNA methylation (**Supplementary** Figures 7-8), and gene length **(Supplementary** Figures 7 and 9), given that these features have been implicated in age-associated transcriptional drift [8,9].

Lee and colleagues reported that genes with CpG islands (CGI+) are less likely to change with age than genes without CpG islands (CGI-) [9]. Accordingly, we found that the proportion of genes with CpG islands located in their promoters increased as a function of our filtering process, suggesting that as we more rigorously select for age-invariant genes, the more prevalent promoter CpG islands become **(Supplementary** Figure 7A). The transcripts themselves were not enriched for greater %CG content, suggesting there is biological specificity of the function of these islands versus an overall increase in CG content in the region **(Supplementary** Figure 9D). We next investigated whether age-invariant genes also showed greater stability in promoter methylation status during in vitro passaging or in vivo aging using reduced-representation bisulfite sequencing (RRBS) datasets. For mouse embryonic fibroblasts serially passaged into senescence, we found both age-variance (based on our skin tissue-specific notation) (**Supplementary** Figure 8A), and CGI (**Supplementary** Figure 8B) status influenced methylation variability. Regardless of age-invariant RG status, CGI+ genes are more stable than CGI-genes **(Supplementary** Figure 8A). However, this pattern was not observed in mouse tissues, including liver, brain, heart, lung, or WBC **(Supplementary** Figure 8B).

Stroeger et al. report that median transcript length is the factor most associated with age-related change, with longer transcripts tending to be downregulated and shorter transcripts tending to be upregulated with age [8]. Complementing these findings, we found that age-invariant genes tend to be shorter than age-variant genes when comparing minimum transcript length(**Supplementary** Figure 7B). However, the opposite is true when comparing either maximum (**Supplementary** Figure 9A) or Ensembl canonical (**Supplementary** Figure 9C) transcript length.

## Discussion

Much of aging biology research has focused on changes that occur across the organismal lifespan and the contribution of these changes to age-related mortality, morbidity, and functional decline [1,38]. Molecular signatures that are robust to aging – specifically, age-invariant genes – have received comparatively little attention. Identifying age-invariant genes allows for further study of why they do not change with age. Lessons from these age-resilient genes provide a complementary view of aging and the stability of biological systems with time. Also, from a practical perspective, because many genes change with age, it is important to identify age-invariant genes for use as **reference genes (RGs)** for gene expression normalization [13]. By adopting a pipeline for identifying RGs from RNA-seq data, we find that there are, in fact, hundreds to thousands of age-invariant genes per tissue. Strikingly, there is poor agreement between the pan-tissue age-invariant genes and commonly used classical RGs. According to our results, none of the classical RGs are suitable for use in cross-sectional aging studies across the 17 tissues studied **(Figure 3A)**, and some canonical tissue-gene pairings (e.g., GAPDH in the liver) are not age-invariant [43]. Our novel age-invariant genes are, therefore, better suited than classical RGs for performing normalization for RT-qPCR experiments in aging tissues.

We report nine pan-tissue age-invariant genes in mice (**Table 1**). Reference and housekeeping gene literature postulates that continuously and stably expressed genes serve essential cellular and organismal functions [12]. Consistent with this hypothesis, depletion of 7 out of 9 of our pan-tissue age-invariant genes have already been reported to induce cell (1110004F10Rik) or embryonic lethality when completely knocked out (Brk1, Rer1, Psmd4, Reco2, Tomm22, and Fis1) [44–49], according to the Mouse Genome Informatics database (www.informatics.jax.org) or International Mouse Phenotyping Consortium database (www.mousephenotype.org). The remaining two transcripts, Srp14 and Gemin7, have no reported knockout mouse strain or phenotypes, but we hypothesize would be lethal if absent.

Two biological processes— mitochondrial function (Fis1, Rexo2, and Tomm22) and proteostasis (Psmd4, Rer1, and Srp14)— emerge from these 9 genes. Although these biological processes are implicated in aging changes, they may also contain components that remain highly stable across the lifespan. Rexo2 (RNA exonuclease 2) was recently shown to increase mitochondrial gene transcription, mediate RNA turnover, and enforce promoter specificity in mammalian mitochondrial transcription [48]. Rer1 returns rogue ER-resident proteins or unassembled subunits in the Golgi apparatus back to the endoplasmic reticulum [46]. Little is known about the molecular function of the small acidic protein 1110004F10Rik (also known as Smap) or its human ortholog C11orf58, but given its high stability and requirement for cell survival, this protein may merit further attention [44]. Thus, the stability of these 9 genes may have evolved as a result of these genes being critical for mitochondrial and proteostatic function, and for continued life in the face of age-related deterioration.

Simply including older mice in our study and utilizing the standard RG identification pipeline was insufficient at filtering out age-invariant genes. Rather, selecting for age-invariant genes required an additional step of explicitly removing genes that are correlated with age. We also find that the variance in expression of a given gene often changes across life stages. For instance, we identified more genes having high variance in young age than in middle or old ages (**Supplementary** Figure 4). Although perhaps surprising, this finding is consistent with reports indicating the proportion of genes decreasing in variance with age is greater than those increasing in variance with age [6,7,50]. It is possible that younger animals show greater variance related to circadian rhythms, the estrous cycles, sex differences, response to stress, or other adaptive and cyclical factors.

Some limitations and caveats constrain our study. First, some of the specific cutoffs we utilized were based on prior work, while others (e.g., exact age correlation cutoff) were based on our best judgment. We provide a complete table of filter results in **Supplementary Table 2** in case others wish to utilize different cutoffs in selecting RGs. To ensure the list of genes provided are useful reference genes in normalization strategies, including RT-qPCR and even some RNA-sequencing normalization approaches, we required high transcript expression through Filter 4. Although consistent with normalization transcript identification strategies in RNA-seq, many low-expression age-invariant genes are absent. Thus, our lists report age-stable, high-expression genes only. Our findings are influenced by the technical limitations of RNA-seq [10,51] and the analytical limitations of high dimensional data, including subsampling of highly heterogeneous samples like aged organisms previously described in the literature [10,51,52]. However, variance in sample collection, processing, and preparation across these datasets likely compensate for any individual source’s batch and degradation bias (e.g., each of the four datasets used employs a different poly-A sample preparation kit). Our final 9 pan-tissue age-invariant genes have been tested individually in 17 tissues and four datasets, totaling 1120 samples, thereby reducing the risk of, for example, a type I error (wrongly identifying a gene as age-invariant). Finally, an important assumption not usually discussed in aging transcriptome literature may influence interpretation in the context of aging: consistent RNA mass. A few studies suggest a decline in total cellular RNA mass with aging [53,54]. This is different from the reported downward trend of differentially expressed genes with age [3]. Current RNA sequencing analysis techniques use proportional estimates (counts per million, fragments per kilobase of transcript per million, transcripts per million, etc.) to normalize samples in order to compare transcript dynamics across samples. Similarly, RT-qPCR protocols typically rely on standardizing total RNA input. If total RNA mass reduction is a global feature of cellular aging, our age-invariant genes are proportionally stable but may decrease in mass with age. Similarly, a gene identified to be overexpressed in old age may maintain constant molar concentration within a cell or tissue. We recommend readers keep these considerations in mind when interpreting any gene expression study in the context of aging.

The existence and study of age-invariant genes have the potential to provide the field of aging with novel insights. It was interesting to find that age-invariant genes were enriched for some pathways associated with hallmarks or pillars of aging **(Figure 4)**, specifically nutrient sensing, proteostasis, mitochondrial function, and immune function. This is somewhat puzzling given that such hallmarks are defined by changes thought to play putatively causal roles in aging [22,55] indeed, genes that most clearly change with age are enriched in the same hallmarks [2]. It is possible enrichment in pathways associated with hallmarks of aging may simply reflect the fact that hallmarks of aging are broad and cover much of biology. In that case, it may be necessary to more specifically delineate each hallmark of aging, e.g., perhaps only a subset of nutrient sensing processes should be considered as a hallmark. However, this broadness would not explain why some hallmarks of aging are associated while others are not. What might be the significance of genes associated with hallmarks of aging that remain stably expressed throughout aging? We note that a prior report indicated that essential genes are enriched for pro-longevity functions, as experimental overexpression of essential genes tends to increase lifespan in yeast [56]. We also find that age-invariant genes are present in pathways linked to human age-related diseases **(Figure 4A-B)**. If age-invariant genes are essential for life, then organisms may have evolved mechanisms to keep these genes stable in the face of pervasive age-related changes in the rest of the pathway or network. One potential example highlighted here is the age-invariant gene enrichment of protein complexes in the electron transport chain. NADH:ubiquinone oxidoreductase, or Mitochondrial Respiratory Complex I, is the only age-invariant gene-enriched ECT complex throughout most tissues **(Figure 4C)**. Although the downregulation of ETC genes is one of the most established transcriptional events in aging [52] and protein Complex I proteins undergo major changes in abundance with age [57], stability in some ETC components is likely required for continued life. This is consistent with Complex I being one of the ETC complexes that can be traced back to the last universal common ancestor of all living organisms [58]. Significant dysregulation of such essential components may be incompatible with life, and evolutionary forces may ensure stability throughout the lifespan. It will be interesting to determine whether further bolstering the expression or stability of such age-invariant genes may be a pro-longevity strategy or, if given their continous expression stable genes are good aging pharmacological targets. The putative aging intervention metformin, for example, may benefit from the stable expression of it’s target, Complex I [59].

In contrast, age-invariant genes were not enriched in some hallmarks, including epigenetic alterations, cellular senescence, and the extracellular matrix. Our results suggest that these three are the most vulnerable to aging as not many genes related to these hallmarks resist age-related change. In agreement with this finding, these hallmarks are key targets across many existing longevity interventions, i.e., epigenetic reprogramming, senolytics, and enhancing extracellular matrix homeostasis [60–62]. Considering that age-invariant genes tend to be essential for life, one hypothesis is that early changes in these hallmarks may not be particularly detrimental for the organism and thus lack the selective pressure to evolve stability mechanisms in aging. The cumulative long-term burden of changes, however, may contribute to pathological aging. Alternatively, these variant hallmarks may reflect adaptive processes that evolved to change dynamically with aging for the benefit of the organism.

Future analyses could focus on the processes that maintain the stability of age-invariant genes. Our initial investigations demonstrate that age-invariant genes are enriched in CpG islands, consistent with a previous report that genes with CpG islands are more resistant to age-related dysregulation than those without CpG islands, which are misexpressed during age-related heterochromatin decondensation [9]. However, further analyses are needed to determine whether the resistance to changes in the methylome of CpG-rich promoters was responsible for the stability of gene expression over time. For instance, whether increased CpG density is better able to reinforce a stable epigenetic state.

We also found that age-invariant genes tend to be shorter than others, confirming a previous study reported that the longest genes show the greatest degree of downregulation [8]. Further study is needed to better understand the relationship between expression dynamics and transcript length. Of note, classical RGs in general have been reported to exhibit shorter introns and exons, low promoter region conservation, 5’ regions with fewer repeated sequences, low nucleosome formation potential, and a higher SINE to LINE ratio [10,24]. It will be important to determine if and how these factors may contribute to the stability of age-invariant genes.

Lastly, it will be important to determine the translatability of our age-invariant transcripts, both to other organisms as well as to protein expression. In a recent study, 52% of human reference genes were matched to independently analyzed mouse reference gene orthologs [14]. Protein abundance can be inferred from transcriptomic data at the tissue and single-cell level, particularly for genes continuously and stably expressed [63,64]. These transcripts show a high correlation (∼0.7) with their protein product except when variability is introduced by cellular state and microenvironment conditions. Given that age-invariant genes are assumed to be expressed in steady-state, many of these genes may also be age-invariant at the protein level.

Here, we provide the aging field with a list of 9 pan-tissue age-invariant genes for use in normalization strategies, e.g., RT-qPCT; we observe that age-invariant genes are enriched in ontology terms associated with some, but not all, hallmarks of aging; and we explore some common features of age-invariant genes (CpG island status and transcript length). Be it for understanding the basic biology of aging, establishing rigorous methodology in the field, investigating the mechanisms promoting age-invariance vs. age-variance, or finding aging therapeutic targets, age-invariant genes are an important area of study.

## Methods

### Data Preparation and Normalization

Four datasets were utilized in this analysis. The Discovery Dataset (GSE132040) consisted of 17 male and female tissues from mice spanning the 4 major life span stages **(Figure 1B)**. 11 of 17 tissues were validated with three datasets of bulk-RNA tissue data from male mice: GSE167665, GSE111164, and GSE141252. Count tables were obtained from GEO and normalized as described below. Sample preparation and alignment can be found in their respective publications [2,4,8]. 5 million counts/sample were set as the count threshold for a sample to be included in normalization and further analysis. In the discovery dataset, hierarchical clustering identified a small number of samples that clustered away from their labeled tissue **(Supplementary** Figure 1A), and examination of tissue-specific markers confirmed they may be mislabeled and, therefore, were removed from analysis **(Supplementary** Figure 1B). The number of samples removed per tissue and lifestage can be seen in **Supplementary Table 11** and those used in the rest of the analysis in **Supplementary Table 10**. GEO accession number, tissue type, and life stage counts can be found in **Supplementary Table 12** for validation datasets. Here, intestine labels refer to samples from both the large and small intestine; and brain to those from both the cerebellum and the frontal cortex.

RNA-seq normalization is essential for proper downstream analysis of datasets. In this study, we identified our genes with two normalization approaches: TPM and TMM. The original reference gene discovery approach described by Eisenberg and Levanon in 2013 [10], utilized RPKM normalized data. Around the same time, conversations about proper data processing produced **Transcript Per Million (TPM)**, an intra-sample normalization method that approximates **relative molar RNA concentration (rmc)** [29]. TPM was only incorporated into this RG identification approach in 2019 [25]. Another major strategy for data normalization techniques involves between-sample normalization. To prevent normalization-based artifacts, and given there is no single best normalization approach, the discovery data was normalized with two different approaches: TPM and **Trimmed Mean of M (TMM)** [30]. TMM, an inter-sample normalization method, generates a normalization factor assuming most genes are not differentially expressed. Therefore, TPM is akin to RT-qPCR due to its similarity with rmc while TMM leverages inter-sample information and is less sensitive to gene outliers. Both performed similarly well at identifying RGs in a recent systematic comparison of normalization methods [26].

TPM normalized data was calculated following the formula:

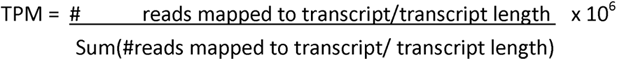

Transcript lengths used in the above formula were obtained with EDASeq package’s (version 3.13) getGeneLengthAndGCContent function. TMM was calculated using the calcNormFactors function from the edgeR package (version 3.40.1).

Gene expression plotting and validation data were performed only with TPM normalized data. Plots were generated with ggplot2(version 3.4.0), ggforce (version 0.4.1) and ggdendro (version 0.1.23).

### Gene Filtering Process

Filters were applied sequentially in R (version 4.2.2) as described in Results. Most mathematical calculations used the r base and MatrixStats package (version 0.63.0). The filter criteria were applied sequentially in both TMM and TPM normalized data, separately for each tissue, thus yielding different lists for each tissue. For each filter, x is either TMM or TPM, and genes were required to pass the filter for both TMM and TPM. Requirements were defined as follows:

1. For each gene: no empty or 0 values
2. For each gene: V, (_2_()) < 1
3. For each gene: V, | _2_()-(_2_()) | < 2
4. For each gene: V, (_2_()) > (_2_()
5. For each gene: %CV≤ 20. V, (_2_()) / (_2_()) 100 < 20
6. For each gene: No correlation with age, based on Pearson’s correlation p-value= 0.05/n. WGCNA package (version 1.71) function corAndPvalue was used to obtain correlation coefficients and p-values. Because each tissue had a 5% chance of finding an association by chance with a fixed 0.05 p-value, a gene present in 17 tissues would have a 58% chance of being erroneously discarded 1-(0.095)^17^. We applied a fractional threshold of a 0.05 p-value, where the p-value threshold applied was 0.05/n, where n is the number of tissues in which the gene in question passed filters 1-4.
7. For each gene: %CV≤ 20 and Spearman correlation p-value= 0.05/n in a validation dataset. n= number of tissues a given gene is present in at filter criteria 6. This step was applied only to TPM normalized data

### RNA isolation and cDNA synthesis

Frozen liver and heart tissues were gifts from Prof. Ron Korstanje at The Jackson Laboratories. Groups consisted of 3 samples per age (8 and 18 months) and sex (female and male), except there was only one sample for an 18-month-old female liver. RNA was isolated with RNeasy Plus Mini Kit (Qiagen #74134) with pestle and syringe homogenization. cDNA was generated using Iscript gDNA Clear cDNA Synthesis (Bio-Rad #1725035) and equivalent RNA mass per 20uL reaction (500ng of heart and 1ug of liver). RNA concentrations were determined with a Qubit 4 fluorometer (Thermo Fisher #Q33238) and RNA BR Assay Kit (Thermo Fisher Q10210).

### Expression data and RG stability

RT-qPCR reactions were assembled with equivalent SsoAdvanced Universal SYBR Green Supermix (Bio-Rad #1725272), cDNA, and respective PrimePCR SYBR Green primers (Bio-Rad #10025636, AssayIDs Atp6v1f: qMmuCID0014923, Cdkn1a:qMmuCED0046265, Srp14: qMmuCID0020464, Tbp:qMmuCID0040542, Tfrc:qMmuCID0039655, Tomm22: qMmuCED0046631). RT-qPCR was performed in a CFX96 thermocycler (Bio-Rad). Stability algorithms NormFinder [35], BestKeeper [33], geNorm [34], and delta-CT method [36] were calculated and integrated into RefFinder [37]. All calculations were performed in R. geNorm and BestKeeper were calculated with the ctrlGene package (version 1.0.1) [65], Normfinder algorithm was downloaded from moma.dk, delta-CT method and RefFinder functions were recreated as originally described. Metadata for the samples used can be found in **Supplementary Table 13**, cycle threshold results in **Supplementary Table 14** for the heart, and **Supplementary Table 15** for the liver.

### CpG island and methylation variability analysis

Gene CpG island (CGI) status was mapped to the annotated list from Lee et al. [9]. Gene names passing each criterion/filter for each tissue were annotated, and percent positive and negative CGI proportion was calculated. Mean and standard deviation were calculated across tissues for each criterion/filter. Counts and percentages of CGI distributions in tissue lists by filter, the odds ratio, statistical test used, and associated p-value are listed in **Supplementary Table 17**.

Composite multi-tissue murine RRBS data [66] was mapped to the mm9 gtf gencode genome. For mouse embryonic fibroblasts, data alignment was previously described [67]. For both datasets, CpG sites common to at least 10 samples and covered by more than 5 reads were analyzed. The methylation status of the promoter region was estimated by averaging the CpG beta values enclosed within 1kb of the transcription start site. Standard deviation was calculated for the methylation of each promoter.

### Enrichment gene analysis

Enrichment analysis was performed using gprofiler2’s (0.2.1) gost function. Electronically annotated GO terms were included in the analysis, and a common custom background of genes expressed at least once in every tissue was imputed. Bonferroni correction was used to calculate enrichment significance. Aging hallmark trajectory enrichment terms were obtained from Schaum et al. [2], while GO biological process terms associated with age-related disease and aging hallmarks were obtained from Fraser et al. 2022 [39]. A few GO terms identified by Schaum et. al. have been discontinued and are marked as obsolete. These terms were excluded from our analysis. Lastly, the top 20 age-invariant GO (biological process, cellular component, and molecular function), KEGG, and Reactome terms were determined by ranking p-values within tissues and taking the lowest 20 gene rank sums across tissues.

For the enrichment maps, all 17 sets of enrichment terms (one per tissue) were used in EnrichmentMap in Cytoscape to generate a consensus network. Different consensus parameters used were used for the CORUM [42] (P-value: 0.05, FDR Q-value: 0.05, Jaccard Overlap Combined: 0.375, test used: Jaccard Overlap Combined Index, k constant = 0.5) and GO:BP terms (P-value: 0.01, FDR Q-value: 0.01, Jaccard: 0.25, test used: Jaccard Index) networks. AutoAnnotate identified common terms for clusters of interconnected nodes. Each node is a pie chart with each slice colored by the enrichment score of each tissue [68].

## Acknowledgements

The research presented here would not have been possible without the exceptional insight of our Biorad Sales Manager, Julie Brunelle. We also thank Profs. Silvia Vilarinho, Rachel Perry and Chen Liu for their insight, support, and advice. We thank Raghav Seghal, Yaroslav Markov, Jessica Kasamoto and Jenel Fraij Armstrong for comments on the manuscript.

## Author Contributions

JTG conceived the study. ATCH, MEL, and JTG designed the study and interpreted the data. JTG performed all experiments and data adquisition in this paper. JTG, MM, and KT developed the R code used. JTG and ATCH wrote the article with feedback from the other authors. All authors approved of the submitted manuscript.

