## Supplementary figures and images for "Age-Invariant Genes: Multi-Tissue Identification and Characterization of Murine Reference Genes"

### Supplemental Figure 1

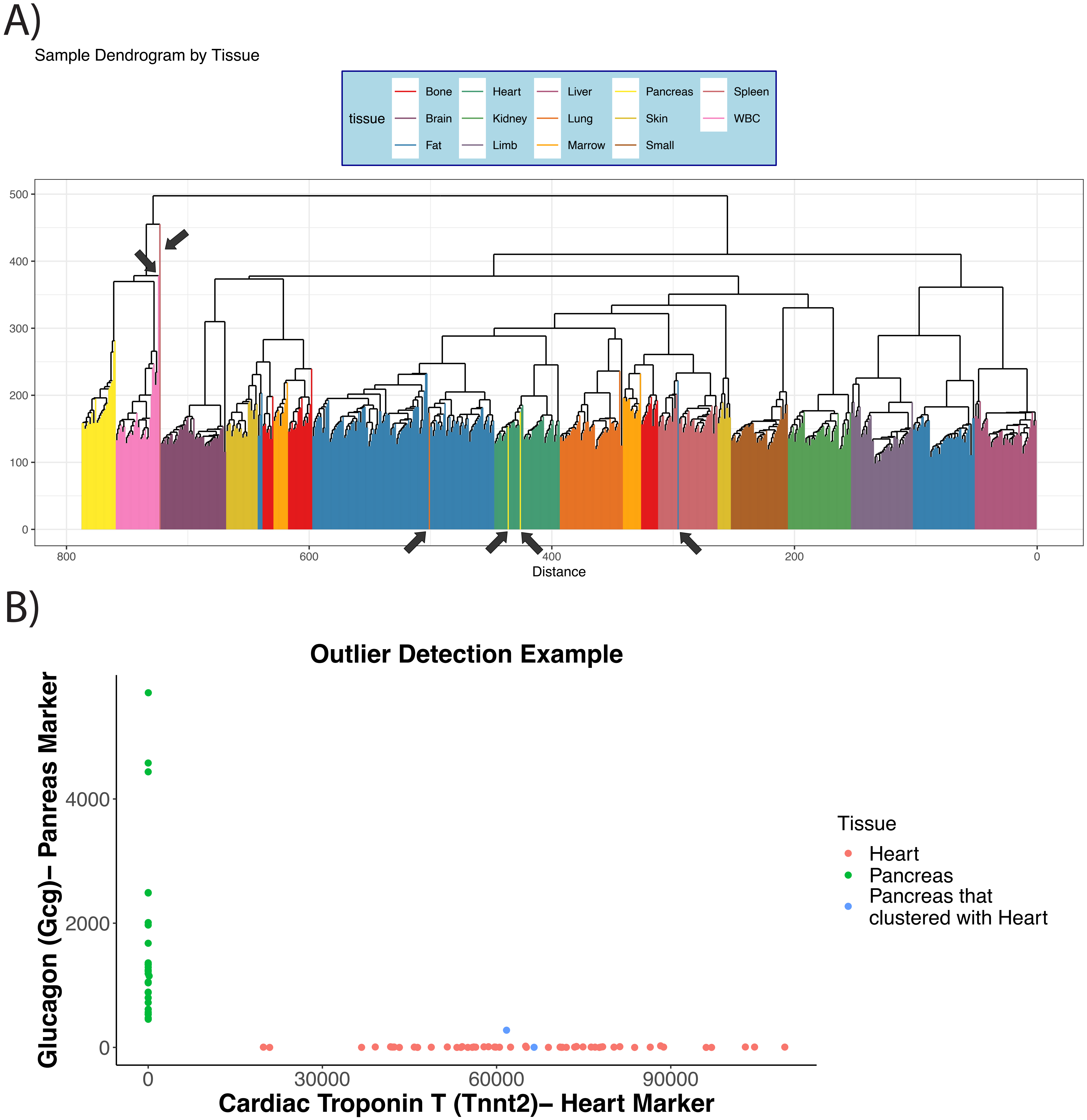

### Supplemental Figure 2

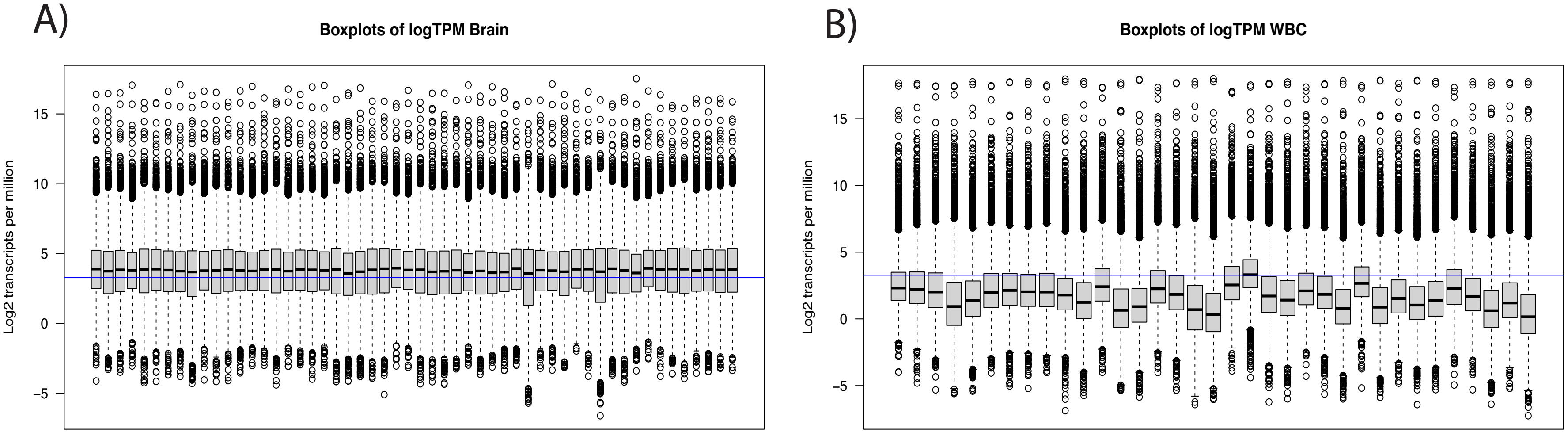

### Supplemental Figure 3

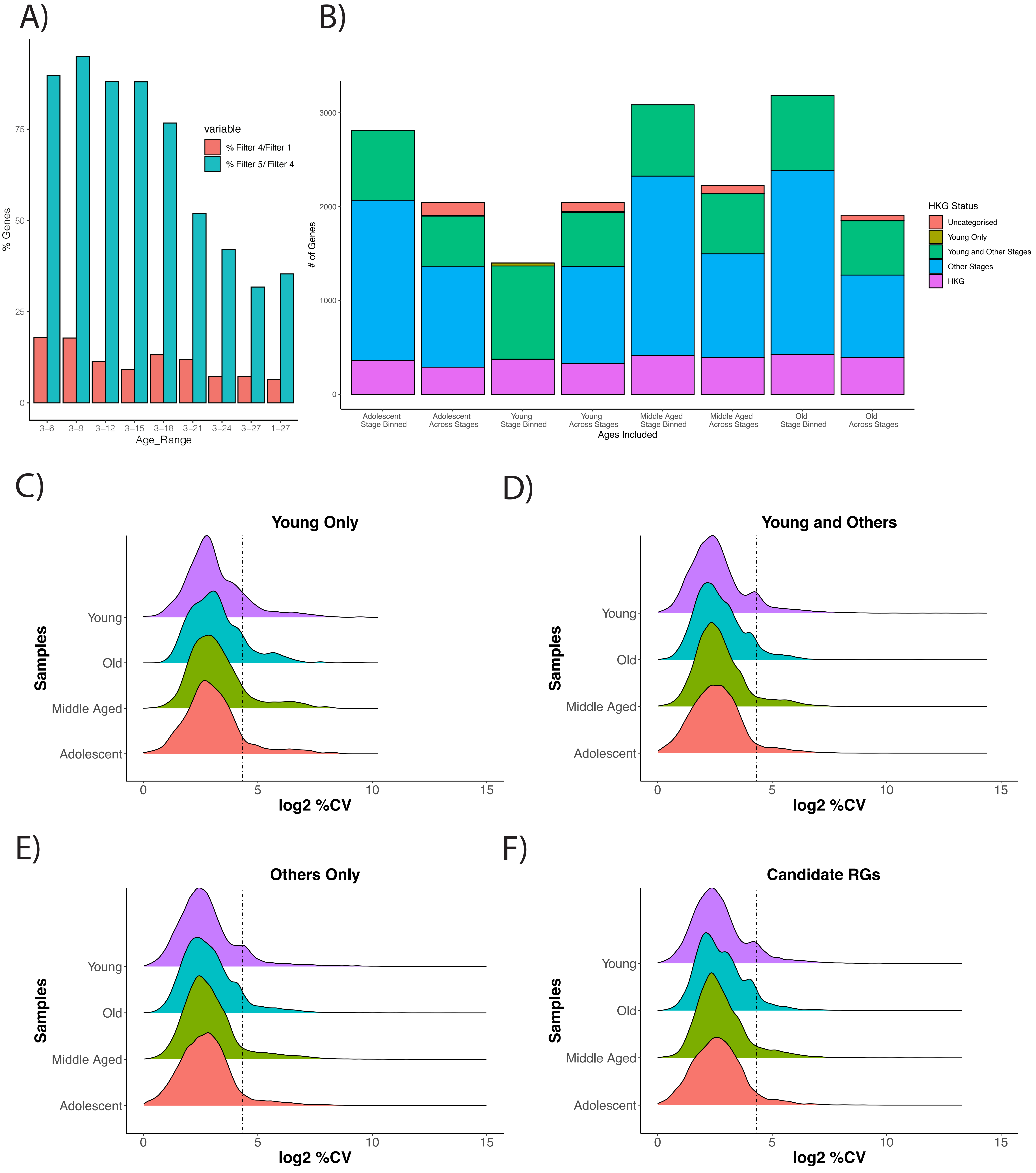

### Supplemental Figure 4

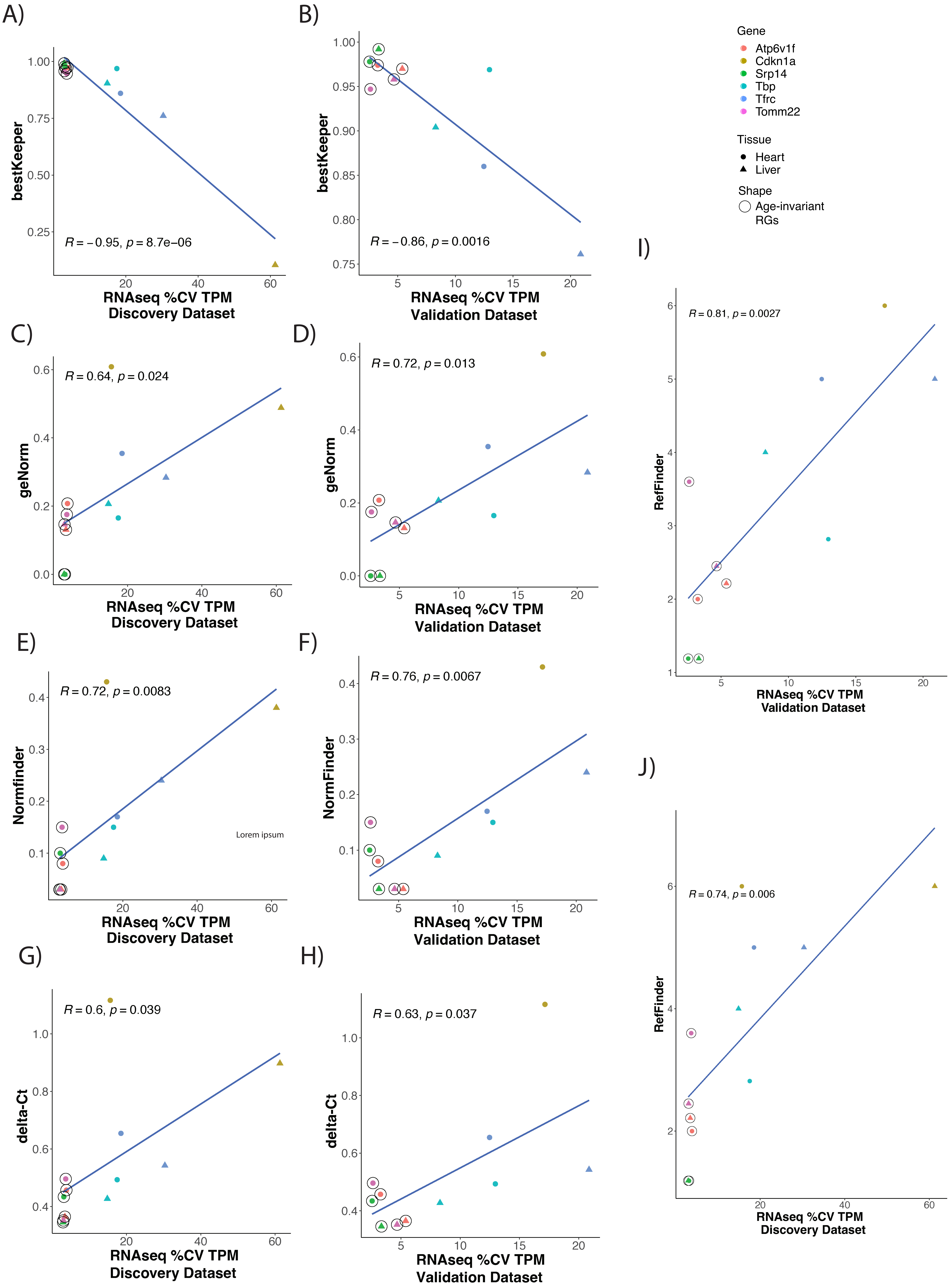

### Supplemental Figure 5

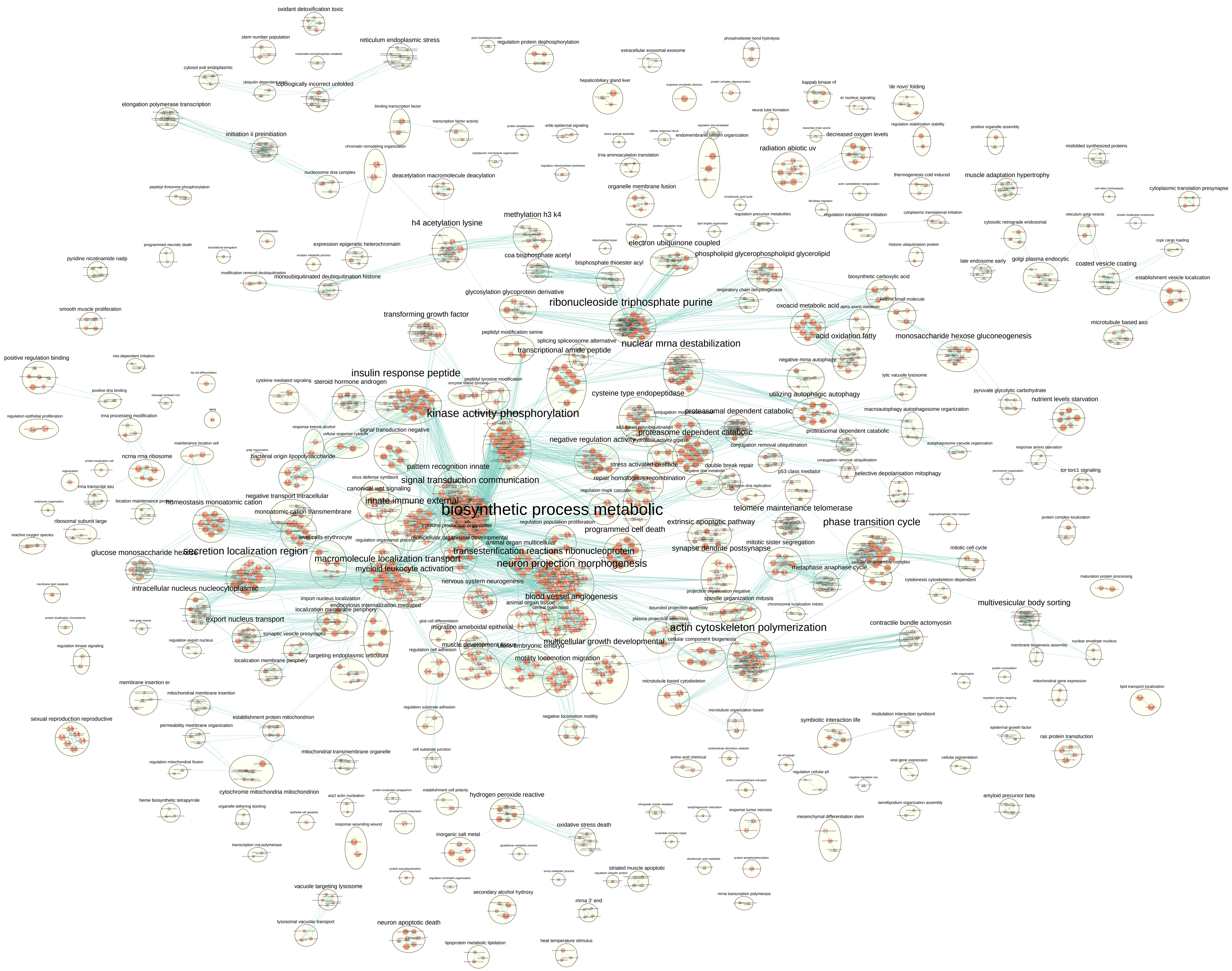

### Supplemental Figure 6

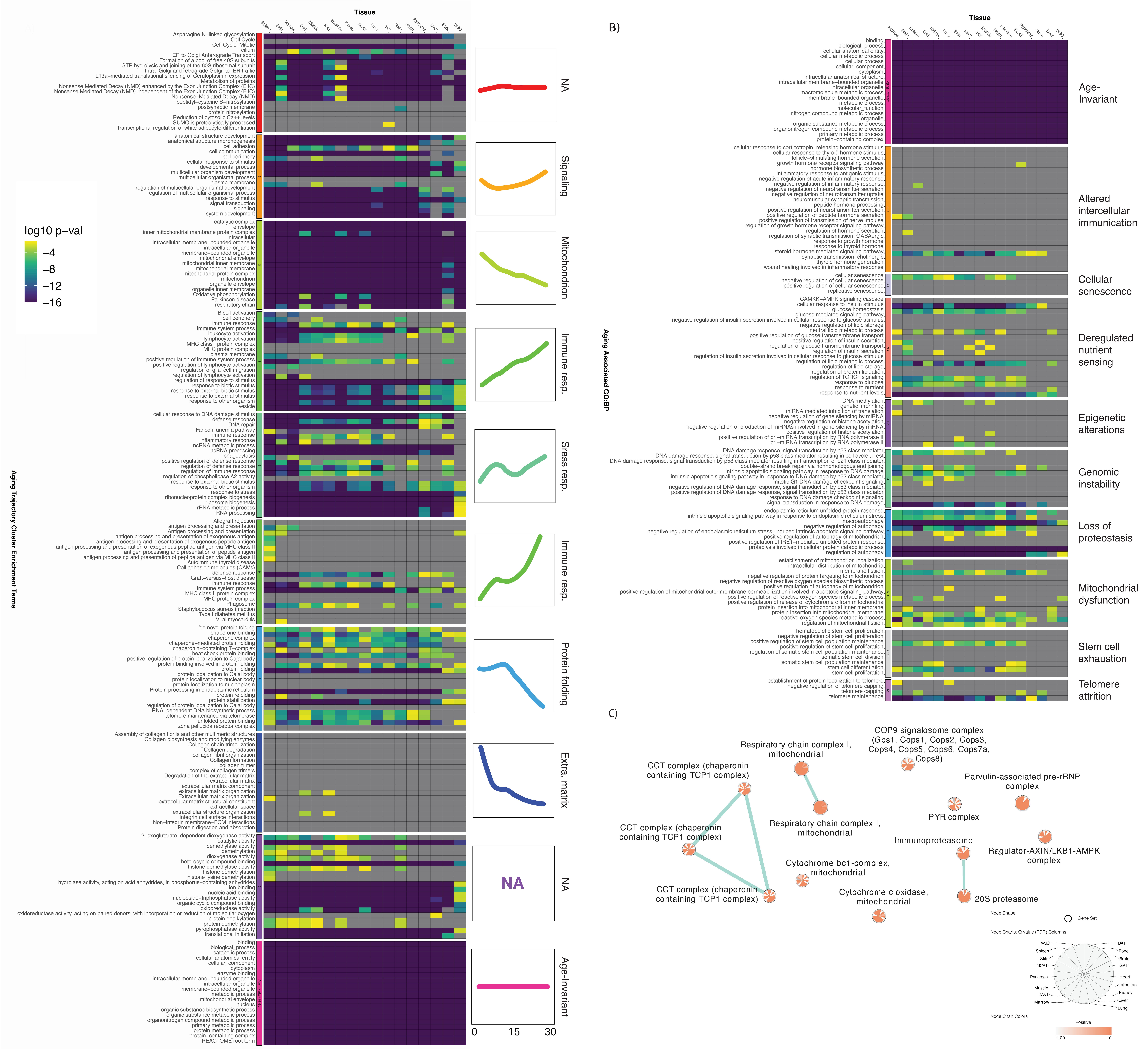

### Supplemental Figure 7

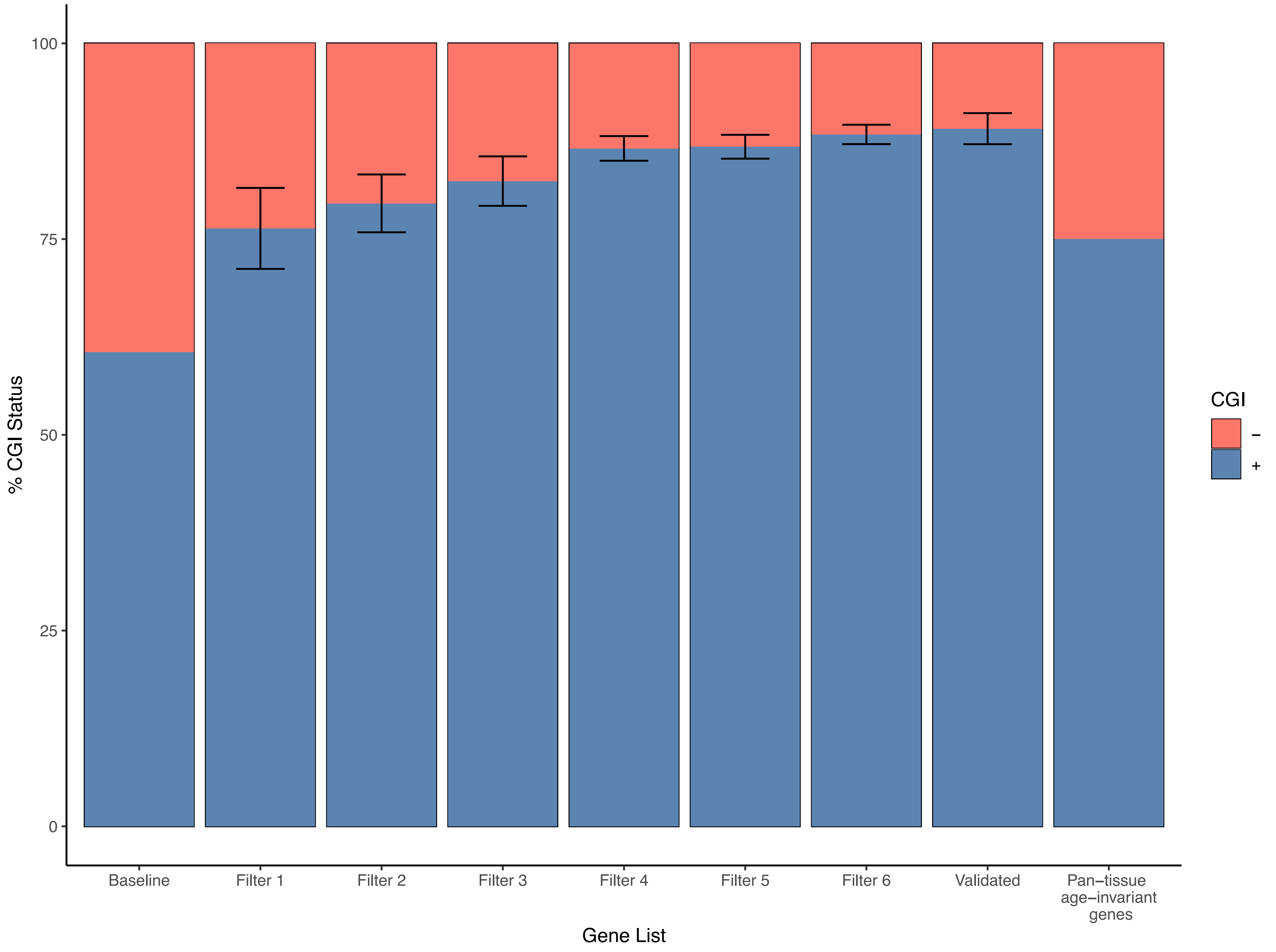

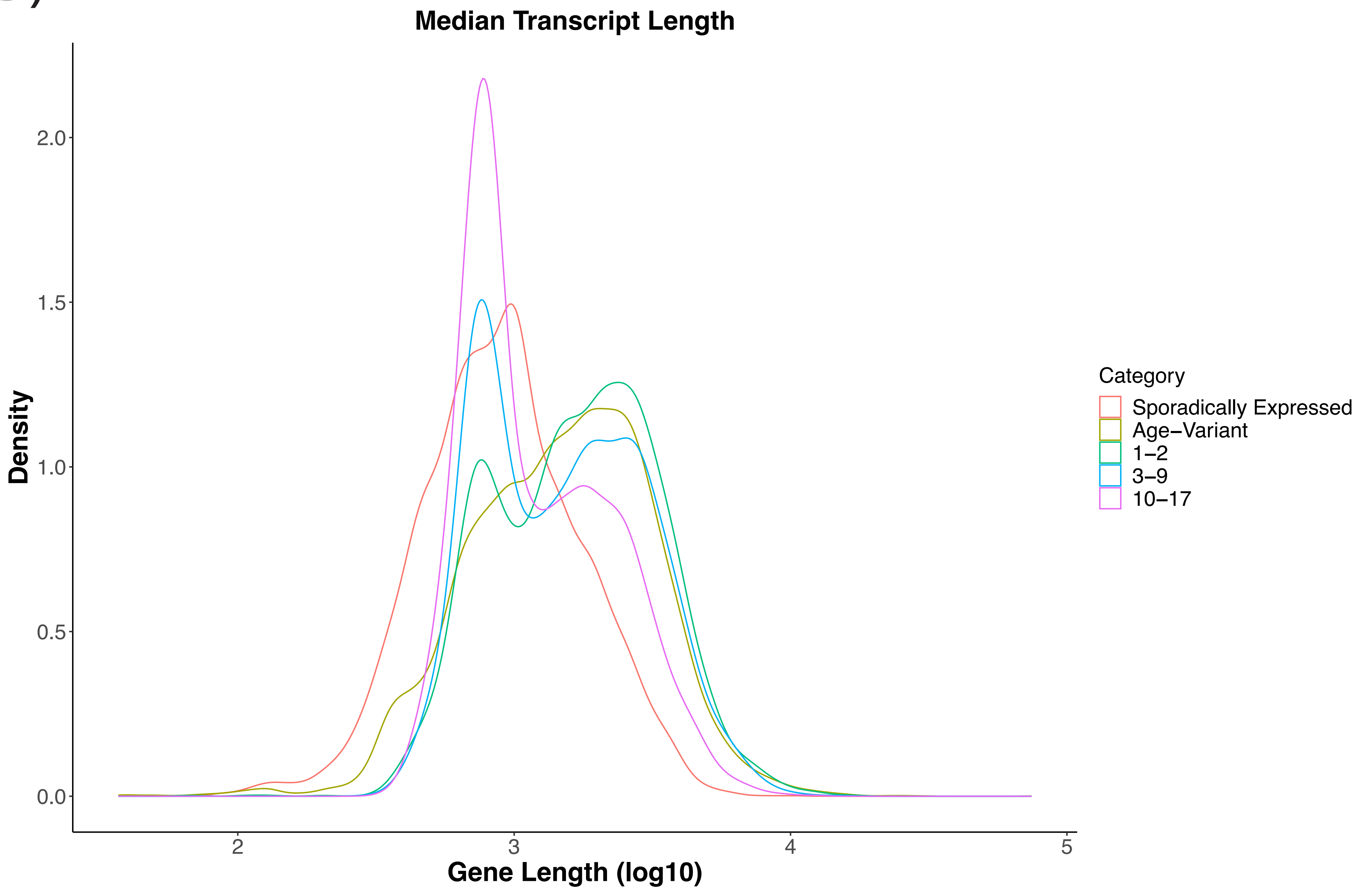

### Supplemental Figure 8

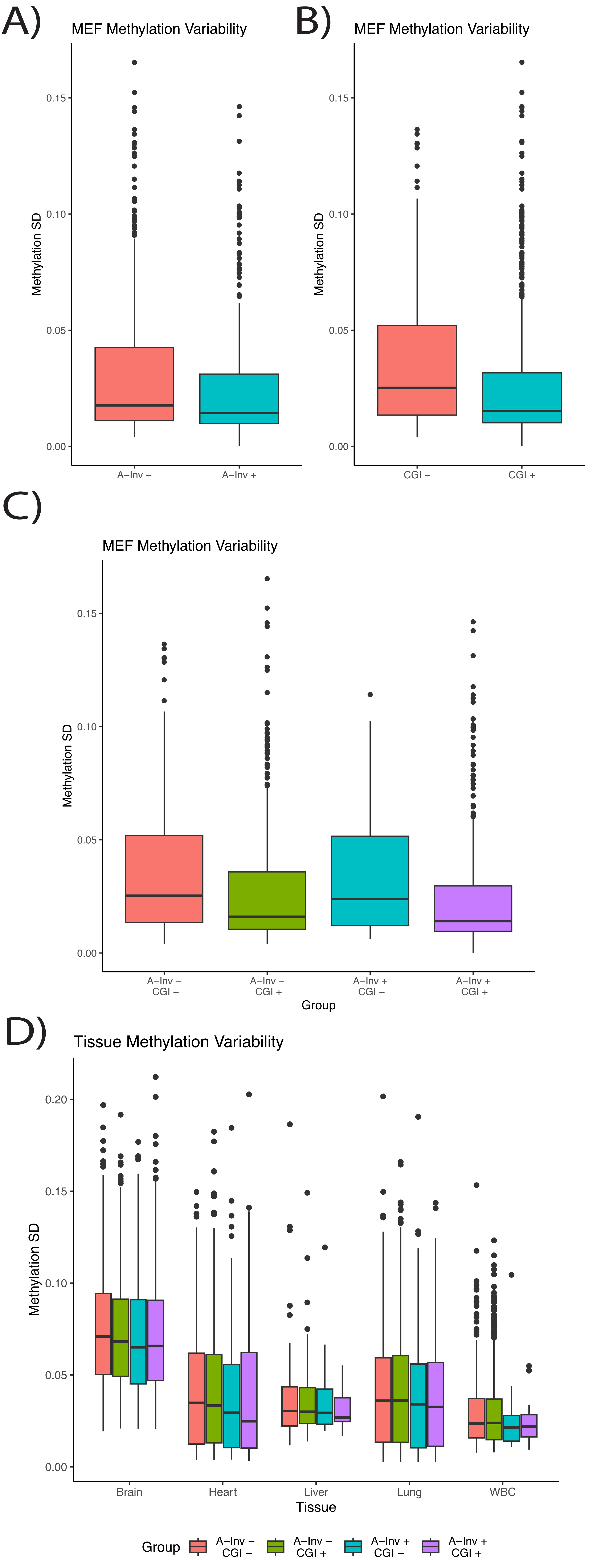

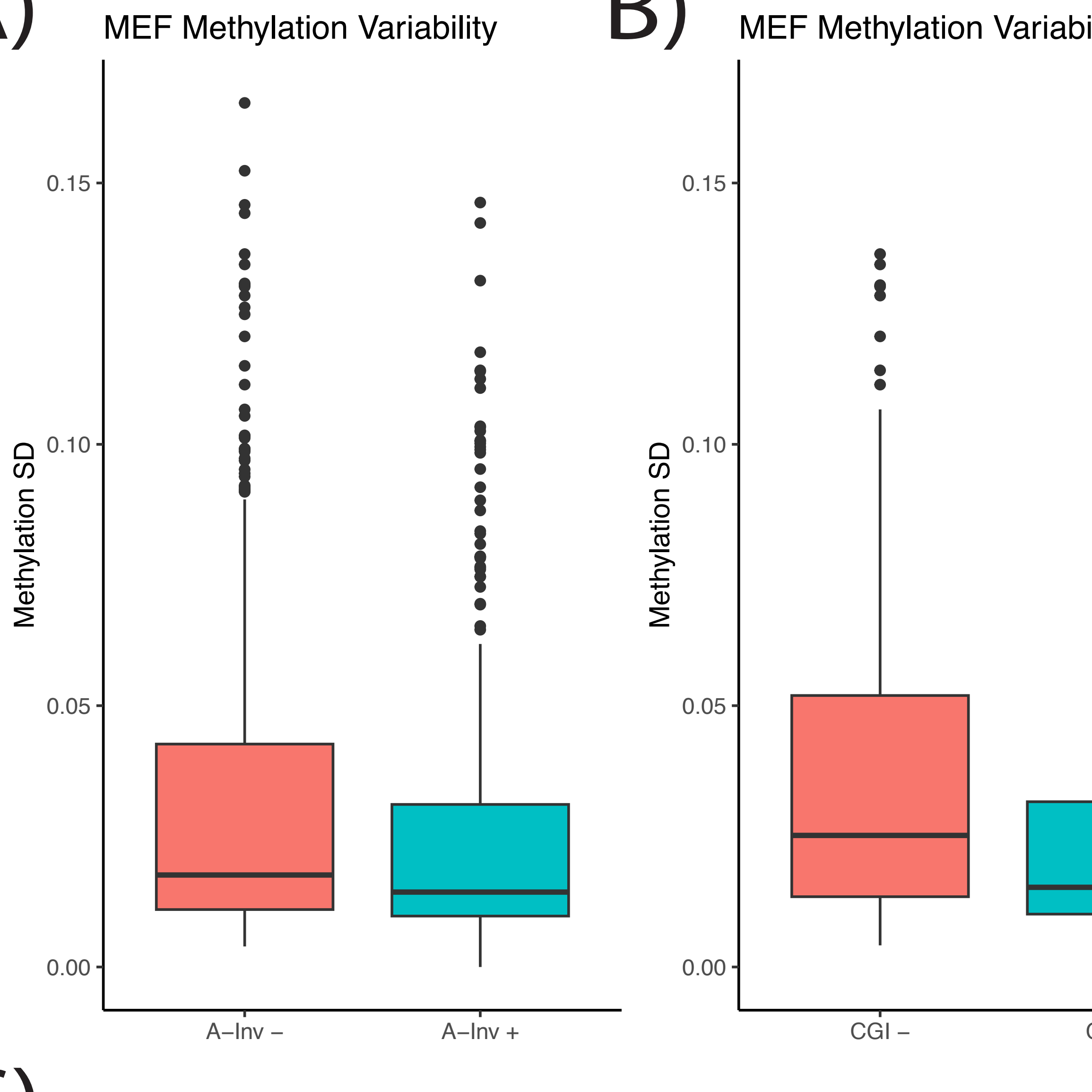

### Supplemental Figure 9

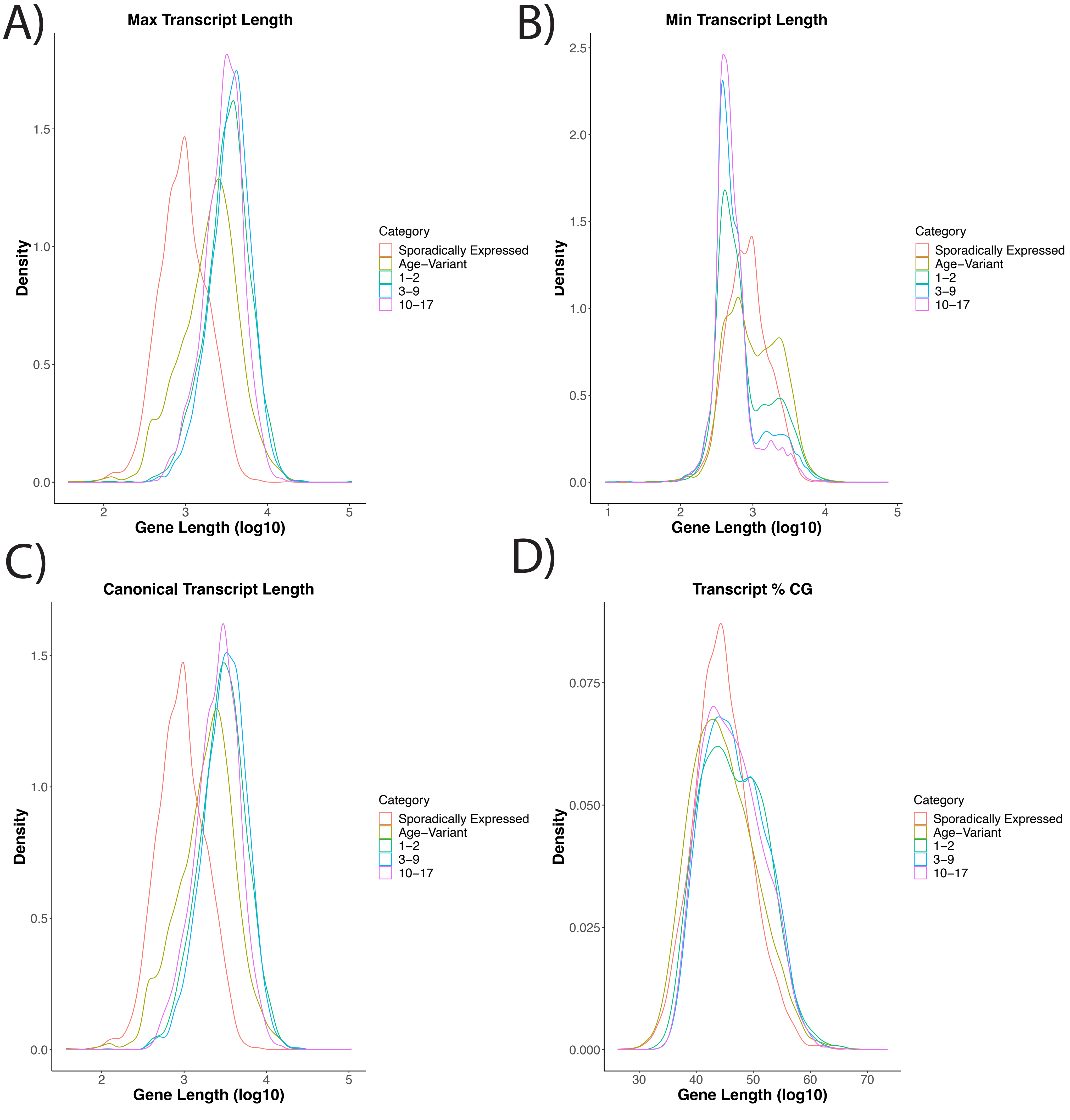
